# Design of post-translational, ligand-controlled inverters and switches

**DOI:** 10.64898/2025.12.05.692672

**Authors:** Samuel D. Swift, Kieran S. McCarthy, Zachary T. Baumer, Ian Wheeldon, Sean R. Cutler, Timothy A. Whitehead

**Affiliations:** Department of Chemical and Biological Engineering, University of Colorado, Boulder; Boulder, 80305, USA; Department of Chemical and Environmental Engineering, University of California, Riverside; Riverside, CA, 92521, USA; Center for Industrial Biotechnology, University of California, Riverside; Riverside, CA, 92521, USA; Institute for Integrative Genome Biology, University of California, Riverside; Riverside, CA, 92521, USA; Department of Botany and Plant Sciences, University of California, Riverside; Riverside, CA, 92521, USA

**Keywords:** Biosensing, Post-translational Signaling, Inverter, Switch, Ultrasensitivity, Modeling, Protein Engineering, Chemically Induced Dimerization, Engineered Living Cells

## Abstract

Rational control over diverse, ligand-responsive output networks is a foundational challenge in synthetic biology, particularly for systems based on post-translational signaling. Here, we present the design and engineering of minimal, modular protein architectures for chemically responsive molecular inverters and digital switches. Our inverter design modifies the plant-derived PYR1-HAB1 chemically inducible dimerization module by incorporating a constitutive activator, which is then competitively displaced by the ligand-bound PYR1 receptor, converting the native ‘dimerization-on’ mechanism into a ‘signal-off’ inverter. We establish key design features and demonstrate predictable tuning of the inverter’s transfer function, including its maximum output, half-maximal inhibitory concentration, and minimum output, solely by adjusting protein stoichiometry. The architecture is modular, enabling plug-and-play response to diverse, user-defined drug-like small molecules. We show that an inverter biosensor for an environmental contaminant functions in engineered living cells with a low nanomolar sensitivity. Additionally, we convert the PYR1-HAB1 sensor into a digital switch by adding an engineered molecular titrant to the system. Overall, this work provides a generalizable, minimal, and tunable protein scaffold for programming complex, post-translational signaling logic, significantly expanding the toolkit for sophisticated biological circuit design.

## Introduction

Transducing a small molecule input(s) into a physiological response is foundational to biological systems and involved in processes as diverse as vertebrate development^1^, plant cell fate and development decisions ^2,3^, perception of odors ^4^, and cell to cell communication in bacteria ^5^. Such responses can be digital (ultrasensitive or all or none) ^6,7^, or analog signals that turn on or off a signal in response to a controlling ligand. The establishment of diverse, ligand-responsive output networks is a fundamental aim in synthetic biology.

Transcriptional and translational circuits have been used to control gene expression via genetically encoded feedback loops^8–12^ to engineer diverse output modalities. These tools have been used to engineer switches^13^, oscillators^14^, and diverse logic gates^15,16^. However, such systems typically process information on the order of hours ^17,18^. By comparison, post-translational circuits enable near-instantaneous sensing because their signal transduction mechanisms are not intertwined within genetically encoded feedback loops, and are therefore more attractive options for certain sensing and therapeutic applications^19–22^. Currently, there are two ways to design such post-translational circuits. First, bespoke proteins can be designed for each novel ligand-receptor pair^19,20^. Alternatively, one can identify mutations which can cause signal inversions or bandgap behavior in ligand-binding proteins; such mutations have been observed in proteins as diverse as plant hormone receptors ^23^, quorum sensing receptors ^24^, and bacterial allosteric transcription factors ^25^. However, both of these strategies require extensive protein engineering and thus remain experimentally intensive and difficult. What is needed is an alternative strategy enabling ligand-modular protein sensing architectures to be output-modular.

In this study, we design two modular post-translational transfer functions (an inverter and a switch) for chemically induced dimerization (CID) modules. We derive biochemical network models that govern the behavior of the inverter and the switch, map their functional space, and present strategies for each that can be used to precisely tune their transfer functions to user defined outputs. We integrate these systems into the PYR1-HAB1 CID module, a biosensing platform whose malleable binding pocket enables facile redesign for diverse small molecules ^26–28^. We validate both the inverter and the switch with a split-luciferase protein complementation plate assay^29^, extend the inverter results to biosensors for several other ligands, and demonstrate the system *in vivo*. All together, in this work we redesign the already *ligand-modular* PYR1-HAB1 platform to also be *output-modular*, broadening its portability and utility.

## Results

### Design and validation of a modular, chemically responsive inverter

We define a chemically responsive inverter as a system of proteins that turns off an output signal in the presence of a controlling small molecule. To design such a system, we chose to modify natural CID modules. CID modules^30^ typically have a ‘dumbell’ type dose-response curve ^31^ where an intermediate concentration of ligand leads to a high amount of complex; low and high concentrations result in low complex formation. The plant-derived PYR1-HAB1 CID module ^2,32^ is a special type of CID module where a chemical input leads to stable, saturable complex formation ^33,34^ (**Fig 1A**). PYR1 is a receptor that naturally senses the plant hormone abscisic acid (ABA). HAB1 is the CID partner protein which is selectively recognized by the ABA-bound PYR1 receptor.

**Figure 1.**
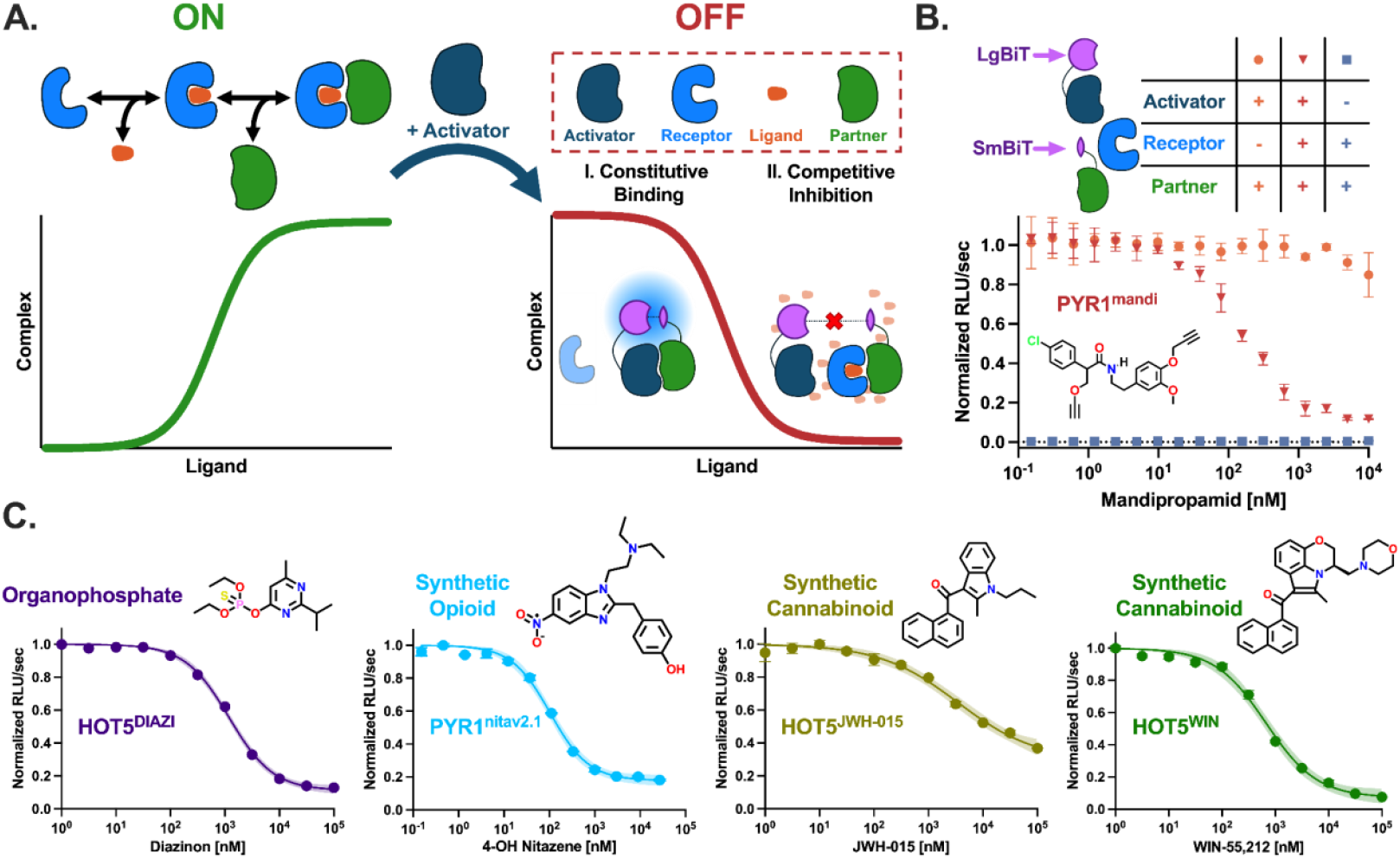
Design of a chemically responsive inverter and validation in diverse sensor-ligand backgrounds. **A**. The left panel shows a cartoon representation of a CID module involving ligand, receptor, and partner. This simple architecture results in a dose-dependent, saturable complex formation. The right panel shows a cartoon representation of a molecular inverter with addition of an activator. Upon ligand addition, the receptor outcompetes the activator using a competitive displacement mechanism. Purple shapes represent the SmBiT and LgBiT pieces of the Nanoluc luciferase; luminescence is used as a proxy for complex formation. **B**. Normalized relative luminescence units (RLU) per second vs. mandipropamid concentrations. Orange circles represent LgBiT-activator and SmBiT-partner. Blue squares represent the receptor (PYR1^mandi^) and SmBiT-partner. Red inverted triangles represent experimental conditions with all three proteins. **C**. The inverter system is portable to biosensors sensitive to diverse ligands. The inhibitor and ligand used is shown above each dose-response curve. Experimental data were fit with a nonlinear least-squares regression to a four-parameter logistical sigmoidal model. A 95% confidence interval of this fit is shown in addition as a shaded region. Data represent the mean (n = 3) and error bars represent the standard error of the mean, although in some cases the error bars are smaller than the symbols themselves.

We desired the minimal components that would turn the PYR1-HAB1 molecular ratchet into an inverter. To this end, we investigated the addition of an activator protein. In our system, an activator protein binds the HAB1 partner protein constitutively (ligand-independent); dimerization of the two elicits a proximity-based output signal (**Fig 1A**). Addition of a receptor - here, an engineered PYR1 - can competitively displace the activator with increasing amounts of a controlling ligand. The ligand-bound receptor outcompetes activator for binding to partner, resulting in loss of the output signal by competitive displacement. The PYR1 receptor has been engineered to bind a wide palette of controlling drug-like ligands^26,27,29,35,36^ with specificity and high affinity, without modifying the HAB1 partner protein or the activator. Therefore, our architecture results in modular ‘plug and play’ inverters for diverse, user-defined drug-like small molecules.

We formulated a biochemical network model for this system (**Equations 1-13**) and identified the key design features that maximizes inhibition (**Figure S1**). First, the concentration of the receptor in the system should exceed that of the partner protein; full inhibition generally requires a 10-fold excess of receptor. Also, the inhibition portion of phase space is enriched where the receptor exceeds the activator. Second, we find inhibition is maximized when the binding affinity between the activator and partner is weaker than that for the ligand-bound receptor and partner. This differential affinity allows the receptor to competitively displace the activator for partner protein. We used these design features to select the parameters of our system.

The partner protein for the inverter is a computationally stabilized HAB1 (ΔN-HAB1^T+^) ^27^. To select the activator, we screened constitutive proteins identified in a previous engineering campaign ^37^. Many mutations to PYR1 increase its constitutive binding to HAB1 ^38^. We identified an activator (PYR1 S16A, Q24E, D26G, S29Q, H34L, E43D, E68K, Q69E, N70G, D80E, N90S, E102D, T118R, R134K) which bound partner protein with high nM/low *μ*M affinity. This basal affinity is much weaker than the ligand-bound complexes for previously engineered sensors ^27,29,39^ and so is suitable as an activator, according to our model.

We validated the inverter using a previously described *in vitro* luminescence assay ^29^. The Large BiT (LgBiT) and Small BiT (SmBiT) of the NanoBiT complementation reporter system ^40^ were genetically fused to the activator and partner proteins, respectively (**Fig 1B**). For the receptor we used a thermally stabilized PYR1 (HOT5^mandi^) ^28,41^ for the fungicide mandipropamid. In the absence of a receptor, luminescence is low and nearly constant over the entire range of mandipropamid concentrations. Additionally, the absence of activator results in minimal luminescence at all mandipropamid concentrations. Inclusion of all three proteins led to a mandipropamid-dependent signal inversion with an half-maximal inhibitory concentration (IC_50_) of 153 nM (95% c.i. 134-177) (**Fig 1B**). To demonstrate the modularity of this inverter, we performed the same experiment except with four different engineered receptors individually recognizing the organophosphate diazinon, a synthetic opioid, and two different synthetic cannabinoids^27,29,41^. In all four cases, identical experimental conditions led to dose-dependent luminescence decreases **(Fig. 1C)** with nM to low *μ*M IC_50_s depending on the exact sensor. Thus, using our minimal architecture, inverters can be constructed to respond to user-defined drug-like molecules in a plug and play fashion.

### Predictive tuning of the max response, IC_50_, and dynamic range of chemically responsive inverters

The characteristics of an inverter can be quantified by a maximum complex formation (C_max_), the minimum complex formation (C_min_), and IC_50_ (**Fig 2**). Our inverter model predicts that changing certain system parameters can tune the transfer function. For example, the model predicts that C_max_ scales proportionally to the concentration of the partner protein when partner is limiting **(Fig. 2A)**. To demonstrate this phenomenon empirically, we varied the concentrations of the partner protein in the split luciferase assay. As predicted, C_max_ shifts proportionally in response to increasing partner protein concentrations. No statistically significant differences are observed in C_min_ and IC_50_, consistent with the model (Extra sum-of-squares F-test, P = 0.7043).

**Figure 2.**
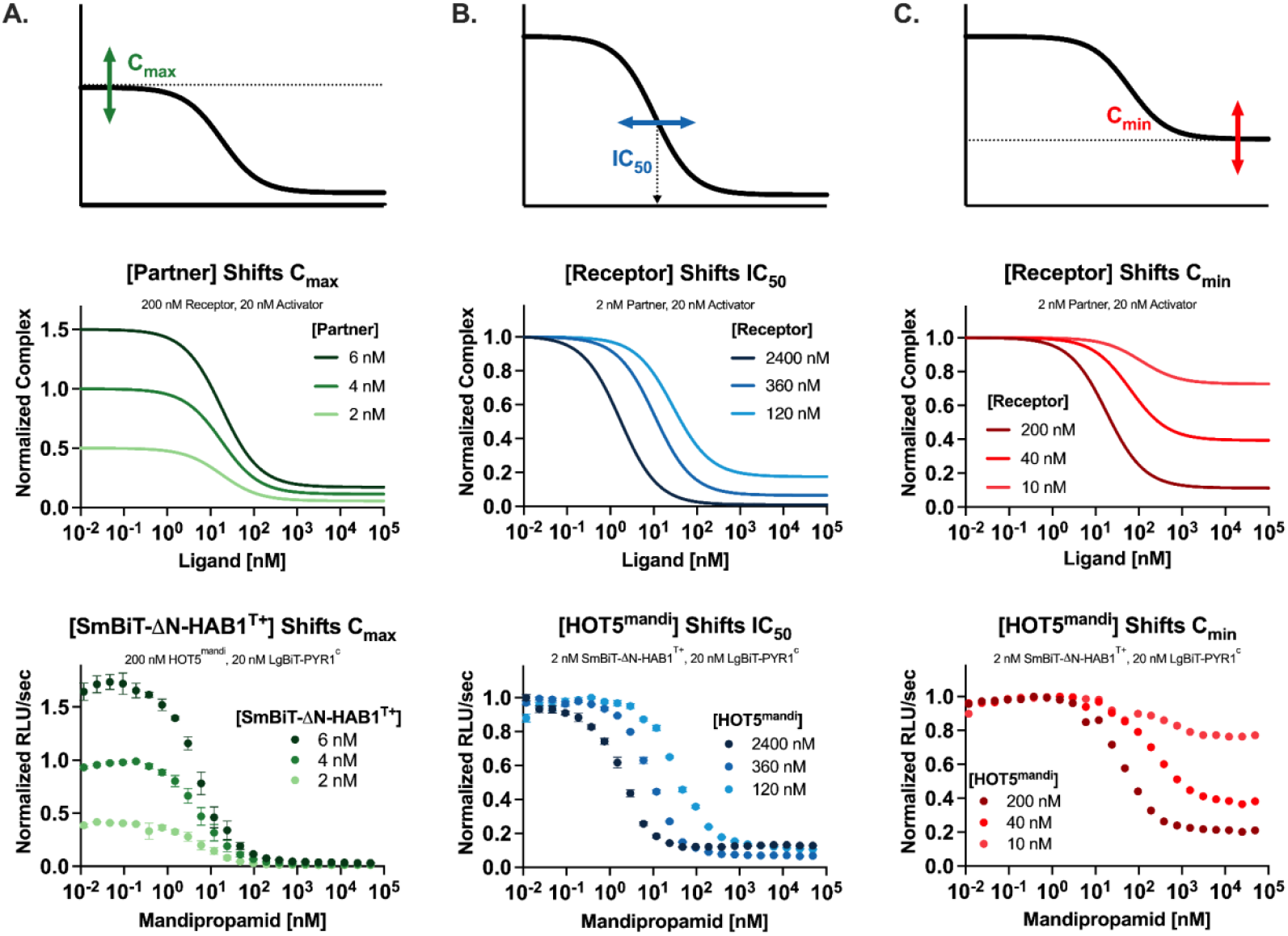
Forward design of the maximum complex formation, half-maximal inhibitory concentration, and minimum complex formation of the inverter. **A**. The maximum complex formation (C_max_) can be tuned predictably by altering the concentration of the partner protein. We modeled this behavior and verified it empirically in an in vitro split-luciferase assay with SmBiT-HAB1 as the partner protein. **B**. The IC_50_ can be tuned by altering the concentration of the inhibitor. We modeled this behavior and verified it empirically in an in vitro split-luciferase assay with HOT5^mandi^ acting as the inhibitor. **C**. The minimum complex formation (C_min_) can be tuned by altering the concentration of the inhibitor. We modeled this behavior and verified it empirically in an in vitro split-luciferase assay with HOT5^mandi^ acting as the inhibitor. The equations used to model the inverter as well as the input parameters used here are included in the methods (**Equations 1-13**). Data represent the mean (n = 3) and error bars represent the standard error of the mean, although in some cases the error bars are smaller than the symbols themselves.

There are several avenues for tuning IC_50_, such as modifying the affinity of the receptor for its ligand. However, most methods require additional protein engineering. Our inverter model predicts tunable control of IC_50_ by changing the ratio of receptor to activator **(Fig. 2B)**. In the presence of a set amount of ligand, increasing the concentration of the receptor also increases the abundance of ligand-bound receptors. More ligand-bound receptors will displace more of the activator, lowering IC_50_. We demonstrate this phenomenon empirically using the split luciferase assay using the mandipropamid receptor. We increased the concentration of the receptor by 3- and 20-fold over baseline, resulting in an IC_50_ shift from 39 nM (95% c.i. 34-45 nM) to 8.8 nM (95% c.i. 8.5-9.2 nM) and 1.9 nM (95% c.i. 1.7-2.1 nM), respectively. Thus, IC_50_ can be predictably tuned for the designed, chemically responsive inverters.

Dynamic range - for an inverter, the extent of inhibition - is governed by C_min_. While C_min_ can be modified using protein engineering, our inverter model predicts C_min_ is tunable via the concentration of the receptor **(Fig. 2C)**. When the receptor is in large fold-excess (>10x) of the activator, % inhibition is at its maximum as more partner protein is competitively displaced under saturating ligand conditions. However, as the ratio of receptor relative to activator is decreased, the fraction of partner protein displaced is decreased–raising C_min_ and decreasing the dynamic range. We demonstrate this phenomenon empirically using the split luciferase assay with the mandipropamid sensor. As we decrease the concentration of the receptor from 200 nM to 40 nM and then 10 nM we observe an increase in C_min_ from 0.21 (95% c.i. 0.19-0.23) to 0.37 (95% c.i. 0.35-0.39) and 0.75 (95% c.i. 0.71-0.79), respectively. Together, the parameters of the inverter transfer function can be predictably tuned guided by a biochemical network model.

#### Engineered living cells signal upon removal of an environmental pollutant

Engineered living cells can be low cost, point of detection biosensors ^42–44^ to inform on contaminants in environmental samples. Mandipropamid is an agriculturally important fungicide used to control late blight in *Solanaceae* like tomatoes, among other applications. However, mandipropamid persists in the watershed of lakes and rivers with half-lives of ∼1 week to ∼1 month respectively ^45^. While the environmental concentrations of mandipropamid fall below levels necessary to be classified as an unacceptable risk for humans^46,47^, they still pose threats to aquatic and terrestrial ecosystems. As such, persistent detection mechanisms are necessary to monitor its environmental concentration. Additionally, mandipropamid is a good proxy for other biocontaminants that necessitate monitoring.

A common problem of engineered living cells as biosensors is the toxicity and/or cellular interference at high ligand concentrations ^48^. An inverter architecture signals upon removal of the pollutant, which improves sensitivity. To demonstrate our inverter *in vivo*, we chose *E. coli* for its extensive genetic toolkit^49^, membrane permeability^44^, and compatibility with luminescence-based assays^40^. Genes encoding the three proteins comprising the inverter system were cloned into a custom plasmid. We sought to control expression ratios between the proteins. To accomplish this, the three open reading frames were expressed as a single mRNA driven by a T7 promoter (**Fig. 3A**). We co-transformed a plasmid encoding an indole-responsive T7 RNA polymerase ^50^ to control the magnitude of protein expression by regulating the amount of mRNA transcribed in an indole-dependent manner (**Fig. 3B**). Additionally, we designed synthetic ribosome binding sites^51,52^ for each gene and incorporated alternative GUG start codons for the activator and partner to decrease their overall expression relative to the mandipropamid receptor ^53^. Finally, we arranged the open reading frames for each protein in order of descending desired concentration^54^.

**Figure 3.**
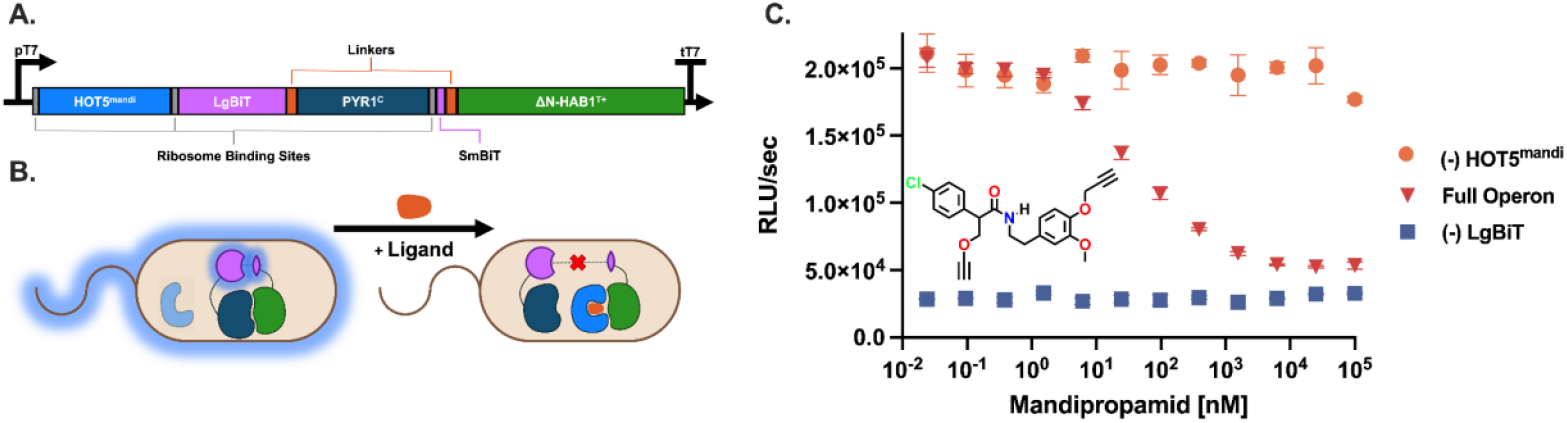
Engineered living cells signal upon removal of an environmental pollutant. **A**. The inverter proteins are expressed as a polycistronic gene under a single T7 promoter. **B**. Schematic of the in vivo inverter within a bacterial host. Indole is used to titer T7 polymerase activity which in turn regulates protein expression levels. In the absence of ligand the cells luminesce which is repressible via ligand induction. **C**. We demonstrate proof-of-concept of the inverter in an in vivo setting using a split-luciferase assay and include relevant controls. Orange circles represent the system with HOT5^mandi^ excised from the operon (n = 2). Inverted red triangles represent the full, intact, inverter operon (n = 4). Blue squares represent the system with LgBiT excised from the operon (n = 2). Data represent the mean and error bars represent the standard error of the mean (n = 4) or the range (n = 2), although in some cases the error bars are smaller than the symbols themselves.

These two plasmids were transformed into *E. coli* and tested in an *in vivo* split-luciferase assay alongside relevant controls (**Fig. 3C**). When the HOT5^mandi^ receptor is excised, no statistically significant differences in luminescence occurred over the entire range of mandipropamid concentrations (horizontal model; Extra sum-of-squares F-test; p-value = 0.08). When the LgBiT is excised, no ligand dependent luminescence is observed. The complete inverter operon yields behavior consistent with its *in vitro* counterpart with an IC_50_ of 39 nM (95% c.i. 31-49 nM). Together, this demonstrates the efficacy of post-translational, chemically responsive inverters in engineered living cells.

### Design and validation of a switch

Ultrasensitivity signaling networks convert small fold-changes in input into large fold-changes in output ^55^. This manifests itself as swift transitions of state and causes systems to exhibit near-binary (digital) outputs. This behavior can be integrated into systems through methods including multimerization, multisite phosphorylation, and positive feedback ^56–58^. To design an ultrasensitive transfer function using the PYR1-HAB1 CID module, we investigated the use of molecular titration (**Fig 4A**) ^59^. As opposed to other approaches like multimerization, adding a titrant protein would involve minimal changes to the parental system.

**Figure 4.**
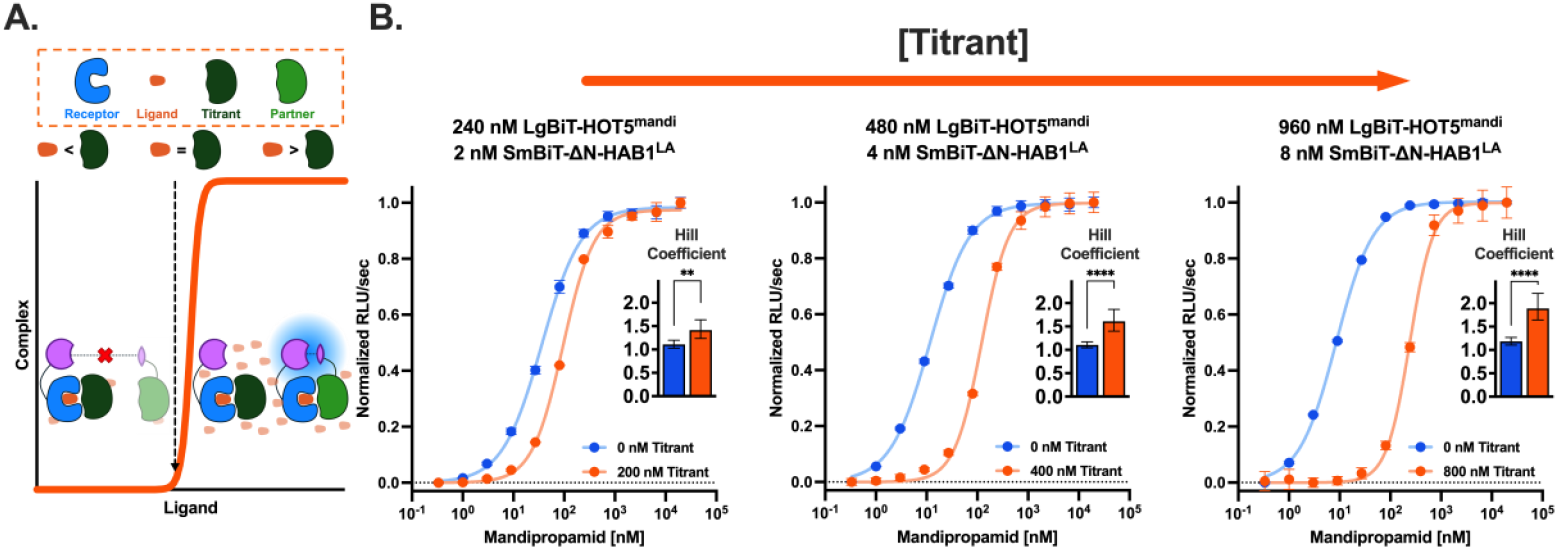
Design and validation of a ligand-responsive switch. **A**. The ligand-responsive switch is facilitated by integrating molecular titration into the PYR1-HAB1 CID module. Here, SpLuc NanoBiT components are included to delineate proteins in the system that constitute functionally active complex. **B**. Increasing the concentration of the titrant, ΔN-HAB1^T+^, titrates the system, represented by higher EC_50s_, and increases the sensitivity of the system, represented by elevated Hill coefficients. Blue circles represent the system without the titrant and orange circles represent the system with. Here, we fit the data with a nonlinear least-squares regression with a specific binding model incorporating a Hill slope. Data represent the mean (n = 3) and error bars represent the standard error of the mean, although in some cases the error bars are smaller than the symbols themselves (Extra sum-of-squares F-Test; **** P < 0.0001, ** P < 0.01).

Molecular titration relies on a protein which sequesters potentially active molecules into inactive complexes^59,60^. For the CID modules, a molecular titrant selectively sequesters the ligand-bound receptor, preventing complexation with the partner protein at low ligand concentrations. Once the number of ligand-bound receptors equals the number of titrant molecules (e.g. the buffer capacity has been reached), any additional ligand added to the system after this point allows the receptor to bind the target protein, triggering an ultrasensitive response.

A closed-form biochemical network model (**Equations 14-24**) shows a narrow operating range of ultrasensitivity for the relative concentrations of the individual proteins and their interwoven affinities **(Figure S2)**. The switch model predicts that sensitivity of the system is maximized when the receptor is in a slight molar excess of the titrant and both proteins are in > 10-fold excess of the partner protein. Additionally, the model predicts that ultrasensitivity requires the ligand-bound receptor to bind the titrant with higher affinity than the partner protein which completes the circuit. Lastly, we model half-maximal effective concentration (EC_50_) and dynamic range as a function of all relative concentrations and binding affinities.

We used ΔN-HAB1^T+^ as the titrant, as it binds engineered ligand-occupied PYR1 receptors at approximately 2 nM ^27,39^. Therefore, we required a HAB1 variant which recognized ligand-bound PYR1 at considerably lower affinities (**<u>L</u>**ow **<u>A</u>**ffinity **<u>H</u>**AB1). We engineered **LAH** (ΔN-HAB1^T+^ E201L, F391K) by screening a focused site saturation mutagenesis HAB1 library using yeast display coupled to fluorescence activated cell sorting **(Fig. S3-S4)**. This library was sorted for diminished binding in the presence of mandipropamid and PYR1^mandi^. This screen yielded several hits. Hits were individually tested and then recombined; we chose **LAH** given its approximate 10-fold reduction in EC_50_ *in vitro* relative to the parental sensor (**Fig. S5)**.

We tested our switch model using a split luciferase assay **(Fig. 4B)**. The LgBiT and SmBiT were fused to HOT5^mandi^ and LAH, respectively. As predicted by the model, as we increase the concentrations of the titrant, ΔN-HAB1, from 200 nM to 400 nM and then 800 nM the Hill coefficients increase from 1.4 (95% c.i. 1.2-1.6) to 1.6 (95% c.i. 1.4-1.9 nM) and then to 1.9 (95% c.i. 1.6-2.2) respectively. Consistent with the model, increasing the titrant also increases the EC_50_, increasing from 94 nM (95% c.i. 85-105 nM) at 200 nM titrant to 241 nM (95% c.i. 222-262 nM) at 800 nM titrant.

## Discussion

Our inverters and switch expand applications of CID modules by enabling new post-translational sensing modalities. The inverter enables repressible activity in an ON-OFF pattern, whereas the switch enables a sharp transition between an OFF and ON state. The novel proteins we engineered for these architectures are ligand agnostic—making the inverter and switch plug-and-play for diverse ligands. We present biochemical network models for both the inverter and the switch, and describe means to tune their transfer functions simply by modifying the stoichiometric ratios of the proteins. This strategy can be readily integrated into CID modules regardless of the parent architecture without the need for bespoke protein engineering.

Fundamentally, the inverter relies on a competitive displacement mechanism to disrupt the basal activity of a functionalized, constitutive PPI with a higher affinity, ligand-sensitive receptor. Because the receptor is independent of the other components, the inverter is readily portable to other ligand-specific sensors - in this work, we demonstrated inverter portability for an agrochemical, an organophosphate, and several synthetic drugs. As modeled, the output signal is infinitely tunable, and the dynamic range can be tuned from a fractional inhibition of 0 to 1. Additionally, we demonstrate a ∼20-fold range of IC_50_ by modifying the concentration of the receptor. The inverter functions *in vivo* while retaining its dynamic range and sensitivity from the *in vitro* results, paving the way for numerous potential applications. For example, engineered living cells with inverters could persistently detect removal of xenobiotics or pollutants in bioremediation efforts. Additionally, inverters could be used to signal upon disappearance of a substrate, which could aid directed evolution of enzymes or for strain engineering.

By implementing molecular titration into the PYR1-HAB1 CID module, we engineer the switch– achieving a Hill coefficient of 1.9. While we only demonstrate the switch for one ligand-sensor, it has been extensively shown that the PYR1-HAB1 platform is portable to diverse ligands^26,29^ and thus we anticipate broad compatibility with additional ligands. We derive a biochemical network model which indicates a narrow functional phase space of the switch; therefore, users should consult these concentration landscapes to guide their desired output and weigh potential tradeoffs between sensitivity, dynamic range, and EC_50_. Consistent with our model, we demonstrate that the Hill coefficient, and EC_50_ of the system are tunable proportional to titrant concentration. However, the extent to which the Hill coefficient increased was relatively lackluster relative to our model’s prediction. The high concentrations of titrant needed to generate ultrasensitive responses have the added defect of introducing proportional levels of constitutive binding of PYR1 and HAB1 ^33^. Constitutive binding is particularly precarious for the switch because it both dampens the dynamic range of the system and muddles the specificity that the system aims to exert, kneecapping immediate translation of the switch for use cases aiming to exploit narrow therapeutic windows or highly specific targets. Together, these limitations impede further translation of the switch into *in vivo* models given the relatively crude tools for controlling protein expression^61^. Potential future directions include protein engineering to remove residual constitutive binding activity between PYR1 and HAB1.

In summary, we have engineered the already *ligand-modular* PYR1-HAB1 platform to also be *output-modular*. While similar biological computing units have been implemented at the transcriptional and translational level, engineering these post-translationally enables rapid sensing and information processing. Synthetic biologists will be able to use the models and underlying techniques outlined here for the chemical control of biology.

## Methods

### Biochemical Network Model of the inverter Transfer Function

The inverter consists of four components: the activator (A), the partner (P), the receptor (R), and ligand (L). The activator binds the partner independent of ligand to form complex (C). Conversely, the receptor must first bind the ligand, forming an intermediate complex (RL), to subsequently form a ternary, sequestered complex (O) with the partner. The elementary rate equations (1-3) along with their corresponding dissociation constant expressions (4-6) that govern the inverter system are shown:

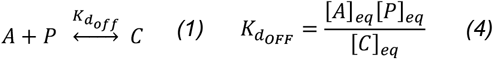

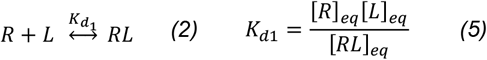

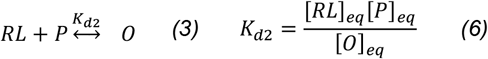

We can then draw upon these governing equations to write net balanced equations (7-10) to relate initial concentrations of each component to a series of equilibrium concentrations:

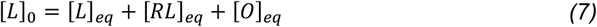

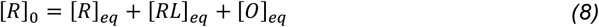

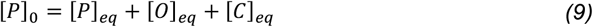

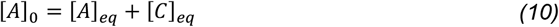

We can then derive a system of equations (11-13) and use bounded non-linear solvers to solve for equilibrium concentrations given initial values of the ligand and proteins.

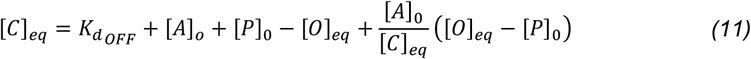

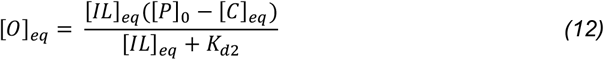

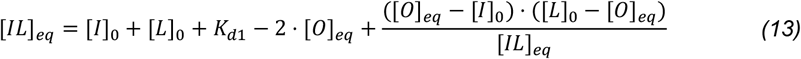

We utilize this model to map out the functional space of the inverter and to inform techniques to tune the output of its transfer function.

### Biochemical Network Model of the Switch Transfer Function

Like the inverter, the switch consists of four components: the receptor (R), the partner protein (P), the titrant (T), and ligand (L). The receptor and ligand bind to form an intermediate complex (RL). The intermediate complex can bind either the titrant to form a sequestered complex (S) or partner protein to form complex (C). The switch transfer function can be modeled as a series of elementary rate equations (14-16) and a corresponding series of dissociation constant expressions (17-19) that govern the system:

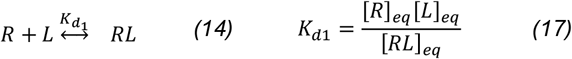

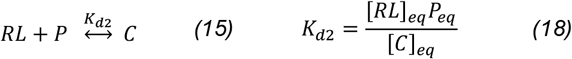

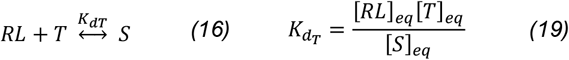

Drawing upon those to derive a series of net balanced equations (20-23) yields:

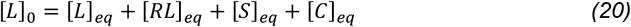

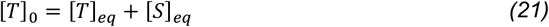

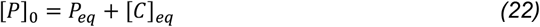

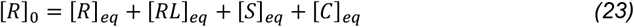

We can then substitute these equations to derive a solution (24) that relates equilibrium complex concentration to all initial concentrations and kinetic constants:

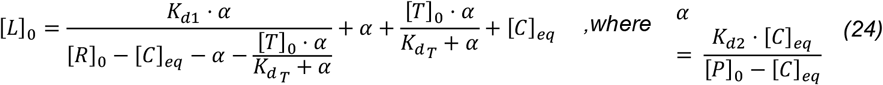

This model is unusual in that a [C]_eq_ is specified to solve for a value of initial ligand [L]_o_ that satisfies the equation. We use this model to map out the functional space of the ultrasensitivity in our system.

### Materials

Mandipropamid (#32805) and diazinon (#45428) were purchased from Sigma-Aldrich. JWH-105 (#10009018), (+)-WIN-55,212-2 (mesylate, #10009023), and 4’-hydroxy nitazene (#30218) were purchased from Cayman Chemical. The luciferin prosubstrate, Hikarazine-108 ^62^, was gifted from Yves Janin at the French National Centre for Scientific Research. All other reagents were sourced from Sigma-Aldrich.

### Plasmid and Library Construction

Plasmids were constructed by HiFi DNA Assembly ^63^, by Golden Gate assembly ^64^, or by site-directed mutagenesis ^65^ using relevant kits from New England Biolabs. A full list of plasmids (**Table S1**), primers (**Table S2**), and synthetic DNA fragments **(Table S3)** are enumerated in the **Supplementary Data**. All primers and synthetic DNA fragments were purchased through Integrated DNA Technologies. All plasmid sequences were verified using whole-plasmid Oxford Nanopore sequencing with Plasmidsaurus.

The HAB1 library was constructed by first using Golden Gate Assembly to insert the ΔN-HAB1^T+^ sequence^27^, coded by ebSDS009, into the yeast surface display (YSD) destination vector, pND003 (Daffern et al.), to create plasmid pSDS101. To create the HAB1 library we used the PyMOL script InterfaceResidues.py to identify twenty-three residues at the HAB1 binding interface of the PYR1^ABA^ and PYR1^mandi^ PDB structures ^28,66^. We downselected eight positions (E201, G246, G247, K381, I383, F391, Y404, L405) that were surface-exposed, not known to play crucial roles in PYR1 binding, and were nonessential for HAB1 folding ^35^. Nicking mutagenesis ^67^ was used for creation of a single site saturation mutagenesis library using primer design script from SUNi mutagenesis ^68^. Deep sequencing showed complete coverage of the 160-member library (**Table S4**). The purified mutagenized plasmid was then transformed into chemically competent EBY100 *S. cerevisiae*, creating the HAB1 YSD library.

### Protein Purification and Preparation

Proteins were expressed in BL21 Star (DE3) and purified via Ni-NTA affinity chromatography as previously described ^27^ with two exceptions. 1. TurboNuclease (AG Scientific, T-1222-50KU) was used in place of Benzonase. 2. MnCl_2_ was omitted in all buffer compositions as we were not utilizing a catalytically active HAB1 variant.

Proteins, stored as ammonium sulfate precipitates, were prepared as described ^29^. Desalting buffer (50 mM HEPES-KOH, 10% w/v glycerol, 200 mM KCl, 1 mM DTT, 5 mM TCEP, pH 8.0) is used for proteins used in the *in vitro* assay, while CBSF++ (20 mM sodium citrate, 147 mM NaCl, 2.5 mM KCl, 0.1% w/v bovine serum albumin, pH 8.0-NaOH) or biotinylation (50 mM HEPES-KOH, 20% w/v glycerol, 200 mM KCl, 5 mM TCEP, pH 8.0) buffers are substituted when applicable.

### Luciferase Plate Assays

*In vitro* assays, involving purified proteins, were conducted exactly as described ^29^. For the *in vivo* experiments, two plasmids, 1) containing the inverter proteins in a polycistronic gene under a T7 promoter (pSDS088, pSDS091, or pSDS092) and 2) containing the LARP-I T7 polymerase ^50^ (pZB578), were transformed into the indole-deficient *E. coli* US0 tnaA-. Single colonies were used to inoculate cultures in SOB supplemented with kanamycin (100 mg/mL) and chloramphenicol (25 mg/mL) and grown overnight for 16 hours at 37°C at 250 rpm. 200 *μ*L of SOB with indole at 400 mM (200 mM working concentration) was added to the bottom of a well in a 96-well, deepwell roundbottom plate (Eppendorf, Cat. No. 951033502). Ligand was diluted to 20% v/v in SOB at a 20x working concentration, and 20 *μ*L were added to the deepwell plate. 180 *μ*L of cultures were then added to the plate to reach a working OD_600_ of 0.4 (final volume 400 *μ*L). This plate was placed on a shaker (Heidolph Titramax/Inkubator 1000) at room temperature (22°C) and 1,050 rpm for an additional 3 hours to induce protein production.

Cells were pipetted up and down to ensure that cultures were well mixed and 100 *μ*L of induced cells were added to a 96-well, white, flat bottom plate (Greiner, Cat. No. 655075). Hikarazine-108 was diluted in PBSF (phosphate buffered saline supplemented with 0.1% w/v bovine serum albumin), to 2x its working concentration of 10 mM. 100 *μ*L of this prosubstrate solution was added to the cell-ligand mixture immediately before running the plate. The plate was placed on a BioTek Synergy™ H1 Microplate Reader and an autogain feature is used at a wavelength of 450 nm, an integration time of 1 second, and a read height of 7 mm. Data was output in units of relative luminescence units (RLU) per second.

### Biotinylation Protocol

Proteins were desalted into biotinylation buffer. 80 molar equivalents of NHS-Biotin (ThermoFisher, Cat. No. 20217) were added to the desalted protein and the solution was incubated for 30 minutes. The reaction was quenched via the addition of Tris-HCl pH 8.0 to 100 mM and DTT to 1 mM and placed on ice until chilled. Ammonium sulfate was added to 80% saturation and the solution was placed on ice for 20 minutes. The precipitated protein was then pelleted via centrifugation at 17,000xg for 10 minutes. At this point, protein can be desalted for immediate use with YSD or resuspended in ammonium sulfate with 5 mM TCEP and 1 mM DTT for long term storage.

### Yeast Surface Display Protocol

All YSD experiments were conducted exactly as described in Steiner et al with the following notable exceptions ^39^. HAB1 was displayed on the yeast surface and PYR1 acted as the biotinylated, labeling protein in the system. Instead of PD-10 columns, Zeba columns (ThermoFisher, Cat. No. 8988) were used to desalt the proteins into CBSF^++^. Representative gating techniques are shown **(Fig. S4)**.

### Engineering <u>L</u>ow <u>A</u>ffinity <u>H</u>AB1

The HAB1 library was chemically transformed into *S. cerevisiae* EBY100 and 10^6^ cells were labeled with 200 nM biotinylated PYR1^mandi^ and 290 nM mandipropamid. The library was gated for displaying, single, yeast cells that had an intermediate level of binding (**Fig. S3C)**. The sorted hits were immediately plated onto SDCAA plates and grown for forty-eight hours at 30°C. Thirty-six colonies were selected, re-induced, and tested at 0 nM, 100 nM, 300 nM, and 900 nM mandipropamid with 200 nM biotinylated PYR1^mandi^. From this screen, four distinct mutations (E201L, F391K, F391A, & F391Y) with moderate loss of binding were identified. Point mutations were tested in the SpLuc *in vitro* assay; mutations showing modest reduction in EC_50_ over the parental background were combined. The E201L and F391K mutations yielded the greatest decrease in PYR1^mandi^ binding affinity. This construct was chosen as the lead LAH candidate and further utilized for proof-of-concept testing of the switch.

### Next-Generation Sequencing Protocol

Quintara AmpExpress was used to deep sequence the HAB1 library. The mutagenized region of HAB1 spans 615 bases therefore the purified library DNA was amplified by two sets of primers, breaking it up into two separate amplicons. FLASH was used to merge pair-end reads ^69^, which were then translated and filtered for quality and length before quantifying mutational frequencies at targeted positions.

### Computational Modeling

Computational models for the inverter system and the switch were implemented using either MatLab, R, or Python. All code is published to: https://github.com/WhiteheadGroup/-Journal-Swift-2026.

## Supporting Information

Concentration phase spaces for the inverter and switch, additional experimental results relevant to engineering LAH (.pdf); tables of plasmids, primers, synthetic DNA, and deep sequencing data (.xlsx).

## Data availability

All data are available in the main text or the supplementary materials. Source Data is provided with this paper. Select plasmids have been deposited in AddGene (deposition #s upon publication).

## Code availability

All code used to generate models for figures is available at https://github.com/WhiteheadGroup/-Journal-Swift-2026. This includes the code used for figures in the SI and to analyze deep sequencing data.

## Author contributions

Conceptualization: SDS, TAW

Methodology: SDS, ZTB, TAW

Investigation: SDS, KSM, TAW

Visualization: SDS, TAW

Funding acquisition: ZTB, SRC, IRW, TAW

Project administration: TAW

Supervision: SRC, IRW, TAW

Writing – original draft: SDS, TAW

Writing – review & editing: SDS, IRW, SRC, TAW

## Acknowledgments

We would like to thank Yves Janin for kindly providing the luciferase prosubstrate, Hikarazine-108.

## Abbreviations

CID: Chemically Induced Dimerization
ABA: Abscisic Acid
LgBiT: Large BiT (NanoBiT Fragment)
SmBiT: Small BiT (NanoBiT Fragment)
C_max_: Maximum Complex Formation
C_min_: Minimum Complex Formation
IC_50_: Half-maximal Inhibitory Concentration
EC_50_: Half-maximal Effective Concentration
RLU: Relative Luminescence Unit

## References

(1) Shimozono, S.; Iimura, T.; Kitaguchi, T.; Higashijima, S.-I.; Miyawaki, A. Visualization of an Endogenous Retinoic Acid Gradient across Embryonic Development. Nature 2013, 496 (7445), 363–366.

(2) Park, S.-Y.; Fung, P.; Nishimura, N.; Jensen, D. R.; Fujii, H.; Zhao, Y.; Lumba, S.; Santiago, J.; Rodrigues, A.; Chow, T.-F. F.; Alfred, S. E.; Bonetta, D.; Finkelstein, R.; Provart, N. J.; Desveaux, D.; Rodriguez, P. L.; McCourt, P.; Zhu, J.-K.; Schroeder, J. I.; Volkman, B. F.; Cutler, S. R. Abscisic Acid Inhibits Type 2C Protein Phosphatases via the PYR/PYL Family of START Proteins. Science. 2009, pp 1068–1071. 10.1126/science.1173041.

(3) Murase, K.; Hirano, Y.; Sun, T.-P.; Hakoshima, T. Gibberellin-Induced DELLA Recognition by the Gibberellin Receptor GID1. Nature 2008, 456 (7221), 459–463.

(4) Buck, L. The Search for Odorant Receptors. Cell 2004, 116, S117–S120.

(5) Waters, C. M.; Bassler, B. L. Quorum Sensing: Cell-to-Cell Communication in Bacteria. Annu. Rev. Cell Dev. Biol. 2005, 21 (1), 319–346.

(6) Ferrell, J. E., Jr; Machleder, E. M. The Biochemical Basis of an All-or-None Cell Fate Switch in Xenopus Oocytes. Science 1998, 280 (5365), 895–898.

(7) Xiong, W.; Ferrell, J. A Positive-Feedback-Based Bistable “memory Module” That Governs a Cell Fate Decision. Nature 2003, 426 (6965), 460–465.

(8) Brophy, J. A. N.; Voigt, C. A. Principles of Genetic Circuit Design. Nat. Methods 2014, 11 (5), 508–520.

(9) Jacob, F.; Monod, J. Genetic Regulatory Mechanisms in the Synthesis of Proteins. J. Mol. Biol. 1961, 3 (3), 318–356.

(10) Winkler, W.; Nahvi, A.; Breaker, R. R. Thiamine Derivatives Bind Messenger RNAs Directly to Regulate Bacterial Gene Expression. Nature 2002, 419 (6910), 952–956.

(11) Bayer, T. S.; Smolke, C. D. Programmable Ligand-Controlled Riboregulators of Eukaryotic Gene Expression. Nat. Biotechnol. 2005, 23 (3), 337–343.

(12) Yen, L.; Svendsen, J.; Lee, J.-S.; Gray, J. T.; Magnier, M.; Baba, T.; D’Amato, R. J.; Mulligan, R. C. Exogenous Control of Mammalian Gene Expression through Modulation of RNA Self-Cleavage. Nature 2004, 431 (7007), 471–476.

(13) Gardner, T. S.; Cantor, C. R.; Collins, J. J. Construction of a Genetic Toggle Switch in Escherichia Coli. Nature 2000, 403 (6767), 339–342.

(14) Elowitz, M. B.; Leibler, S. A Synthetic Oscillatory Network of Transcriptional Regulators. Nature 2000, 403 (6767), 335–338.

(15) Siuti, P.; Yazbek, J.; Lu, T. K. Synthetic Circuits Integrating Logic and Memory in Living Cells. Nat. Biotechnol. 2013, 31 (5), 448–452.

(16) Anderson, J. C.; Voigt, C. A.; Arkin, A. P. Environmental Signal Integration by a Modular AND Gate. Mol. Syst. Biol. 2007, 3 (1), 133.

(17) Yosef, N.; Regev, A. Impulse Control: Temporal Dynamics in Gene Transcription. Cell 2011, 144 (6), 886–896.

(18) Takahashi, M. K.; Chappell, J.; Hayes, C. A.; Sun, Z. Z.; Kim, J.; Singhal, V.; Spring, K. J.; Al-Khabouri, S.; Fall, C. P.; Noireaux, V.; Murray, R. M.; Lucks, J. B. Rapidly Characterizing the Fast Dynamics of RNA Genetic Circuitry with Cell-Free Transcription-Translation (TX-TL) Systems. ACS Synth. Biol. 2015, 4 (5), 503–515.

(19) Shui, S.; Scheller, L.; Correia, B. E. Protein-Based Bandpass Filters for Controlling Cellular Signaling with Chemical Inputs. Nat. Chem. Biol. 2024, 20 (5), 586–593.

(20) Giordano-Attianese, G.; Gainza, P.; Gray-Gaillard, E.; Cribioli, E.; Shui, S.; Kim, S.; Kwak, M.-J.; Vollers, S.; Corria Osorio, A. D. J.; Reichenbach, P.; Bonet, J.; Oh, B.-H.; Irving, M.; Coukos, G.; Correia, B. E. A Computationally Designed Chimeric Antigen Receptor Provides a Small-Molecule Safety Switch for T-Cell Therapy. Nat. Biotechnol. 2020, 38 (4), (1) 426–432.

(21) Olson, E. J.; Tabor, J. J. Post-Translational Tools Expand the Scope of Synthetic Biology. Curr. Opin. Chem. Biol. 2012, 16 (3-4), 300–306.

(22) Mansouri, M.; Ray, P. G.; Franko, N.; Xue, S.; Fussenegger, M. Design of Programmable Post-Translational Switch Control Platform for on-Demand Protein Secretion in Mammalian Cells. Nucleic Acids Res. 2023, 51 (1), e1.

(23) Stammnitz, M. R.; Lehner, B. The Genetic Architecture of an Allosteric Hormone Receptor. bioRxiv, 2025. 10.1101/2025.05.30.656975.

(24) Swem, L. R.; Swem, D. L.; Wingreen, N. S.; Bassler, B. L. Deducing Receptor Signaling Parameters from in Vivo Analysis: LuxN/AI-1 Quorum Sensing in Vibrio Harveyi. Cell 2008, 134 (3), 461–473.

(25) Tack, D. S.; Tonner, P. D.; Pressman, A.; Olson, N. D.; Levy, S. F.; Romantseva, E. F.; Alperovich, N.; Vasilyeva, O.; Ross, D. The Genotype-Phenotype Landscape of an Allosteric Protein. Mol. Syst. Biol. 2021, 17 (3), e10179.

(26) Tian, H.; Beltrán, J.; George, W.; Lenert-Mondou, C.; Seder, N.; Davis, Z. I.; Swift, S. D.; Girke, T.; Whitehead, T. A.; Wheeldon, I.; Cutler, S. R. Unusually Broad-Spectrum Small-Molecule Sensing Using a Single Protein Scaffold. bioRxivorg, 2025. 10.1101/2025.05.15.654352.

(27) Beltrán, J.; Steiner, P. J.; Bedewitz, M.; Wei, S.; Peterson, F. C.; Li, Z.; Hughes, B. E.; Hartley, Z.; Robertson, N. R.; Medina-Cucurella, A. V.; Baumer, Z. T.; Leonard, A. C.; Park, S.-Y.; Volkman, B. F.; Nusinow, D. A.; Zhong, W.; Wheeldon, I.; Cutler, S. R.; Whitehead, T. A. Rapid Biosensor Development Using Plant Hormone Receptors as Reprogrammable Scaffolds. Nat. Biotechnol. 2022, 40 (12), 1855–1861.

(28) Park, S.-Y.; Peterson, F. C.; Mosquna, A.; Yao, J.; Volkman, B. F.; Cutler, S. R. Agrochemical Control of Plant Water Use Using Engineered Abscisic Acid Receptors. Nature 2015, 520 (7548), 545–548.

(29) Leonard, A. C.; Lenert-Mondou, C.; Chayer, R.; Swift, S.; Baumer, Z. T.; Delaney, R.; Friedman, A. J.; Robertson, N. R.; Seder, N.; Wells, J.; Whitmore, L. M.; Cutler, S. R.; Shirts, M. R.; Wheeldon, I.; Whitehead, T. A. Computational Design of Dynamic Biosensors for Emerging Synthetic Opioids. bioRxivorg, 2025, 2025.05.15.654300. 10.1101/2025.05.15.654300.

(30) Stanton, B. Z.; Chory, E. J.; Crabtree, G. R. Chemically Induced Proximity in Biology and Medicine. Science 2018, 359 (6380). 10.1126/science.aao5902.

(31) Cao, S.; Kang, S.; Mao, H.; Yao, J.; Gu, L.; Zheng, N. Defining Molecular Glues with a Dual-Nanobody Cannabidiol Sensor. Nat. Commun. 2022, 13 (1), 815.

(32) Melcher, K.; Ng, L.-M.; Zhou, X. E.; Soon, F.-F.; Xu, Y.; Suino-Powell, K. M.; Park, S.-Y.; Weiner, J. J.; Fujii, H.; Chinnusamy, V.; Kovach, A.; Li, J.; Wang, Y.; Li, J.; Peterson, F. C.; Jensen, D. R.; Yong, E.-L.; Volkman, B. F.; Cutler, S. R.; Zhu, J.-K.; Xu, H. E. A Gate-Latch-Lock Mechanism for Hormone Signalling by Abscisic Acid Receptors. Nature 2009, 462 (7273), 602–608.

(33) Steiner, P. J.; Swift, S. D.; Bedewitz, M.; Wheeldon, I.; Cutler, S. R.; Nusinow, D. A.; Whitehead, T. A. A Closed Form Model for Molecular Ratchet-Type Chemically Induced Dimerization Modules. Biochemistry 2023, 62 (2), 281–291.

(34) Leonard, A. C.; Whitehead, T. A. Design and Engineering of Genetically Encoded Protein Biosensors for Small Molecules. Curr. Opin. Biotechnol. 2022, 78 (102787), 102787.

(35) Park, S.-Y.; Qiu, J.; Wei, S.; Peterson, F. C.; Beltrán, J.; Medina-Cucurella, A. V.; Vaidya, A. S.; Xing, Z.; Volkman, B. F.; Nusinow, D. A.; Whitehead, T. A.; Wheeldon, I.; Cutler, S. R. An Orthogonalized PYR1-Based CID Module with Reprogrammable Ligand-Binding Specificity. Nat. Chem. Biol. 2024, 20 (1), 103–110.

(36) Robertson, N. R.; Lenert-Mondou, C.; Leonard, A. C.; Tafrishi, A.; Carrera, S.; Lee, S.; Aguilar, Y.; Sanchez Zamora, L.; Nguyen, T.; Beltrán, J.; Li, M.; Cutler, S. R.; Whitehead, T. A.; Wheeldon, I. PYR1 Biosensor-Driven Genome-Wide CRISPR Screens for Improved Monoterpene Production in Kluyveromyces Marxianus. ACS Synth. Biol. 2025, 14 (8), 2972–2978.

(37) Baumer, Z. T. Dynamic Regulation of Engineered T7 RNA Polymerases. CU Scholar. https://scholar.colorado.edu/concern/graduate_thesis_or_dissertations/xg94hr36w (accessed 2025-11-17).

(38) Mosquna, A.; Peterson, F. C.; Park, S.-Y.; Lozano-Juste, J.; Volkman, B. F.; Cutler, S. R. Potent and Selective Activation of Abscisic Acid Receptors in Vivo by Mutational Stabilization of Their Agonist-Bound Conformation. Proc. Natl. Acad. Sci. U. S. A. 2011, 108 (51), 20838–20843.

(39) Steiner, P. J.; Bedewitz, M. A.; Medina-Cucurella, A. V.; Cutler, S. R.; Whitehead, T. A. A Yeast Surface Display Platform for Plant Hormone Receptors: Toward Directed Evolution of New Biosensors. AIChE J. 2020, 66 (3). 10.1002/aic.16767.

(40) Dixon, A. S.; Schwinn, M. K.; Hall, M. P.; Zimmerman, K.; Otto, P.; Lubben, T. H.; Butler, B. L.; Binkowski, B. F.; Machleidt, T.; Kirkland, T. A.; Wood, M. G.; Eggers, C. T.; Encell, L. P.; Wood, K. V. NanoLuc Complementation Reporter Optimized for Accurate Measurement of Protein Interactions in Cells. ACS Chem. Biol. 2016, 11 (2), 400–408.

(41) Daffern, N.; Johansson, K. E.; Baumer, Z. T.; Robertson, N. R.; Woojuh, J.; Bedewitz, M. A.; Davis, Z.; Wheeldon, I.; Cutler, S. R.; Lindorff-Larsen, K.; Whitehead, T. A. GMMA Can Stabilize Proteins Across Different Functional Constraints. J. Mol. Biol. 2024, 436 (11), 168586.

(42) Ostrov, N.; Jimenez, M.; Billerbeck, S.; Brisbois, J.; Matragrano, J.; Ager, A.; Cornish, V. W. A Modular Yeast Biosensor for Low-Cost Point-of-Care Pathogen Detection. Sci. Adv. 2017, 3 (6), e1603221.

(43) An, B.; Wang, Y.; Huang, Y.; Wang, X.; Liu, Y.; Xun, D.; Church, G. M.; Dai, Z.; Yi, X.; Tang, T.-C.; Zhong, C. Engineered Living Materials for Sustainability. Chem. Rev. 2023, 123 (5), 2349–2419.

(44) Wu, Y.; Wang, C.-W.; Wang, D.; Wei, N. A Whole-Cell Biosensor for Point-of-Care Detection of Waterborne Bacterial Pathogens. ACS Synth. Biol. 2021, 10 (2), 333–344.

(45) Zhang, J.; Li, Y.; Tan, Y.; Zhang, Y.; Li, R.; Zhou, L.; Wang, M. The Enantioselective Environmental Fate of Mandipropamid in Water-Sediment Microcosms: Distribution, Degradation, Degradation Pathways and Toxicity Assessment. Sci. Total Environ. 2023, 891 (164650), 164650.

(46) Management Board members; Director, E.; Operational Management. Peer review of the pesticide risk assessment of the active substance mandipropamid. European Food Safety Authority. https://www.efsa.europa.eu/en/efsajournal/pub/2935 (accessed 2025-11-21).

(47) Hou, Z.; Hou, X.; Wei, L.; Cao, Z.; Lu, Z.; Liu, H.; Lu, Z. Degradation and Residues of Mandipropamid in Soil and Ginseng and Dietary Risk Assessment in Chinese Culture. Environ. Sci. Pollut. Res. Int. 2023, 30 (10), 26367–26374.

(48) Essington, E. A.; Vezeau, G. E.; Cetnar, D. P.; Grandinette, E.; Bell, T. H.; Salis, H. M. An Autonomous Microbial Sensor Enables Long-Term Detection of TNT Explosive in Natural Soil. Nat. Commun. 2024, 15 (1), 10471.

(49) Ruiz, N.; Silhavy, T. J. How Escherichia Coli Became the Flagship Bacterium of Molecular Biology. J. Bacteriol. 2022, 204 (9), e0023022.

(50) Baumer, Z. T.; Newton, M. S.; Löfstrand, L.; Carpio Paucar, G. N.; Farny, N. G.; Whitehead, T. A. Engineered Stop and Go T7 RNA Polymerases. ACS Synth. Biol. 2024, 13 (12), 4165–4174.

(51) Reis, A. C.; Salis, H. M. An Automated Model Test System for Systematic Development and Improvement of Gene Expression Models. ACS Synth. Biol. 2020, 9 (11), 3145–3156.

(52) Cetnar, D. P.; Salis, H. M. Systematic Quantification of Sequence and Structural Determinants Controlling mRNA Stability in Bacterial Operons. ACS Synth. Biol. 2021, 10 (1) (2), 318–332.

(53) Hecht, A.; Glasgow, J.; Jaschke, P. R.; Bawazer, L. A.; Munson, M. S.; Cochran, J. R.; Endy, D.; Salit, M. Measurements of Translation Initiation from All 64 Codons in E. Coli. Nucleic Acids Res. 2017, 45 (7), 3615–3626.

(54) Svidritskiy, E.; Demo, G.; Korostelev, A. A. Mechanism of Premature Translation Termination on a Sense Codon. J. Biol. Chem. 2018, 293 (32), 12472–12479.

(55) Goldbeter, A.; Koshland, D. E., Jr. An Amplified Sensitivity Arising from Covalent Modification in Biological Systems. Proc. Natl. Acad. Sci. U. S. A. 1981, 78 (11), 6840–6844.

(56) Ferrell, J. E., Jr; Ha, S. H. Ultrasensitivity Part I: Michaelian Responses and Zero-Order Ultrasensitivity. Trends Biochem. Sci. 2014, 39 (10), 496–503.

(57) Ferrell, J. E., Jr; Ha, S. H. Ultrasensitivity Part II: Multisite Phosphorylation, Stoichiometric Inhibitors, and Positive Feedback. Trends Biochem. Sci. 2014, 39 (11), 556–569.

(58) Ferrell, J. E., Jr; Ha, S. H. Ultrasensitivity Part III: Cascades, Bistable Switches, and Oscillators. Trends Biochem. Sci. 2014, 39 (12), 612–618.

(59) Dane Wittrup, K.; Tidor, B.; Hackel, B. J.; Sarkar, C. A. Quantitative Fundamentals of Molecular and Cellular Bioengineering; MIT Press, 2020.

(60) Buchler, N. E.; Louis, M. Molecular Titration and Ultrasensitivity in Regulatory Networks. J. Mol. Biol. 2008, 384 (5), 1106–1119.

(61) Greco, F. V.; Pandi, A.; Erb, T. J.; Grierson, C. S.; Gorochowski, T. E. Harnessing the Central Dogma for Stringent Multi-Level Control of Gene Expression. Nat. Commun. 2021, 12 (1), 1738.

(62) Janin, Y. L.; Coutant, E. P.; Gagnot, G.; Hervin, V.; Baatallah, R.; Goyard, S.; Hibti, F. E.; Quatela, A.; Jacob, Y.; Rose., T. Bioluminescence-Based Reporting Systems of Marine Origin, Few Chemistry Contributions. https://hal.science/hal-03932102v1/preview/Poster%20Y%20Janin%20MBSJ.pdf#page=2.

(63) Gibson, D. G.; Young, L.; Chuang, R.-Y.; Venter, J. C.; Hutchison, C. A., 3rd; Smith, H. O. Enzymatic Assembly of DNA Molecules up to Several Hundred Kilobases. Nat. Methods 2009, 6 (5), 343–345.

(64) Engler, C.; Kandzia, R.; Marillonnet, S. A One Pot, One Step, Precision Cloning Method with High Throughput Capability. PLoS One 2008, 3 (11), e3647.

(65) Zheng, L.; Baumann, U.; Reymond, J.-L. An Efficient One-Step Site-Directed and Site-Saturation Mutagenesis Protocol. Nucleic Acids Res. 2004, 32 (14), e115.

(66) Dupeux, F.; Antoni, R.; Betz, K.; Santiago, J.; Gonzalez-Guzman, M.; Rodriguez, L.; Rubio, S.; Park, S.-Y.; Cutler, S. R.; Rodriguez, P. L.; Márquez, J. A. Modulation of Abscisic Acid Signaling in Vivo by an Engineered Receptor-Insensitive Protein Phosphatase Type 2C Allele. Plant Physiol. 2011, 156 (1), 106–116.

(67) Wrenbeck, E. E.; Klesmith, J. R.; Stapleton, J. A.; Adeniran, A.; Tyo, K. E. J.; Whitehead, T. A. Plasmid-Based One-Pot Saturation Mutagenesis. Nat. Methods 2016, 13 (11), 928–930.

(68) Mighell, T. L.; Toledano, I.; Lehner, B. SUNi Mutagenesis: Scalable and Uniform Nicking for Efficient Generation of Variant Libraries. PLoS One 2023, 18 (7), e0288158.

(69) Magoč, T.; Salzberg, S. L. FLASH: Fast Length Adjustment of Short Reads to Improve Genome Assemblies. Bioinformatics 2011, 27 (21), 2957–2963.

