## Supplementary Information for "Design of post-translational, ligand-controlled inverters and switches"

Affiliations:

**This supplementary material contains:**

Figures S1 to S5

### Supplementary Figures

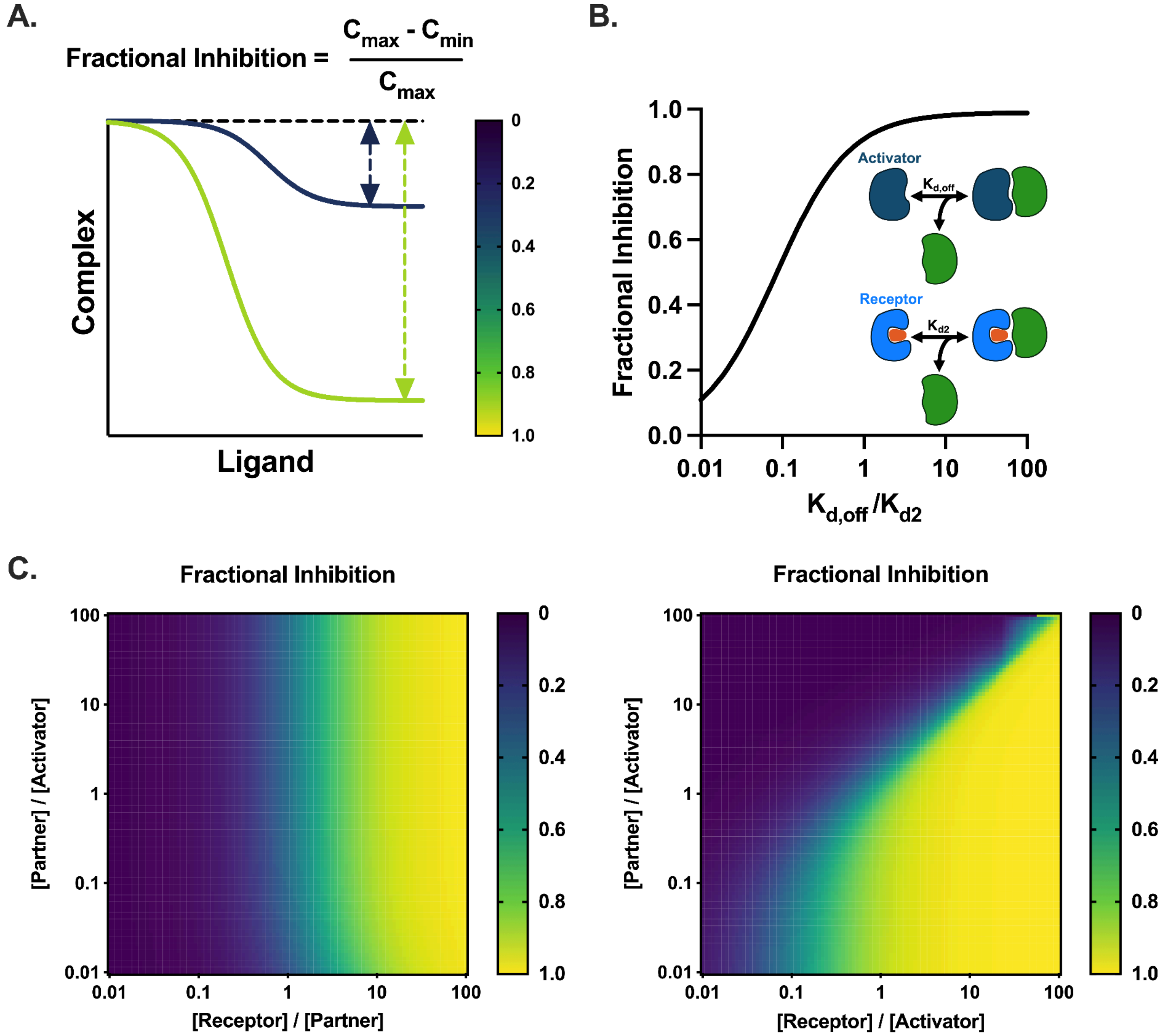


**Figure S1. Optimal phase space of the inverter. A.** The maximum, fractional inhibition of the inverter is defined as the dynamic range over the maximum complex formation. **B.** When partner protein has higher affinity for receptor bound ligand than for activator ($K_{d2} < K_{d,off}$) % inhibition of the inverter system is maximized. Here, we fixed $K_{d2}$ at 2.1 nM and varied $K_{d,off}$; 200 nM receptor, 20 nM activator, 2 nM partner, $K_{d1}$ = 115.6 nM. **C.** When receptor concentration is in >10-fold molar excess of partner protein and a simple molar excess of activator % inhibition of the inverter system is maximized. Here, we set the partner at 2 nM and varied receptor and activator (left) and then set the activator at 20 nM and varied receptor and partner (right); $K_{d2}$ = 2.1 nM, $K_{d,off}$ = 200 nM.

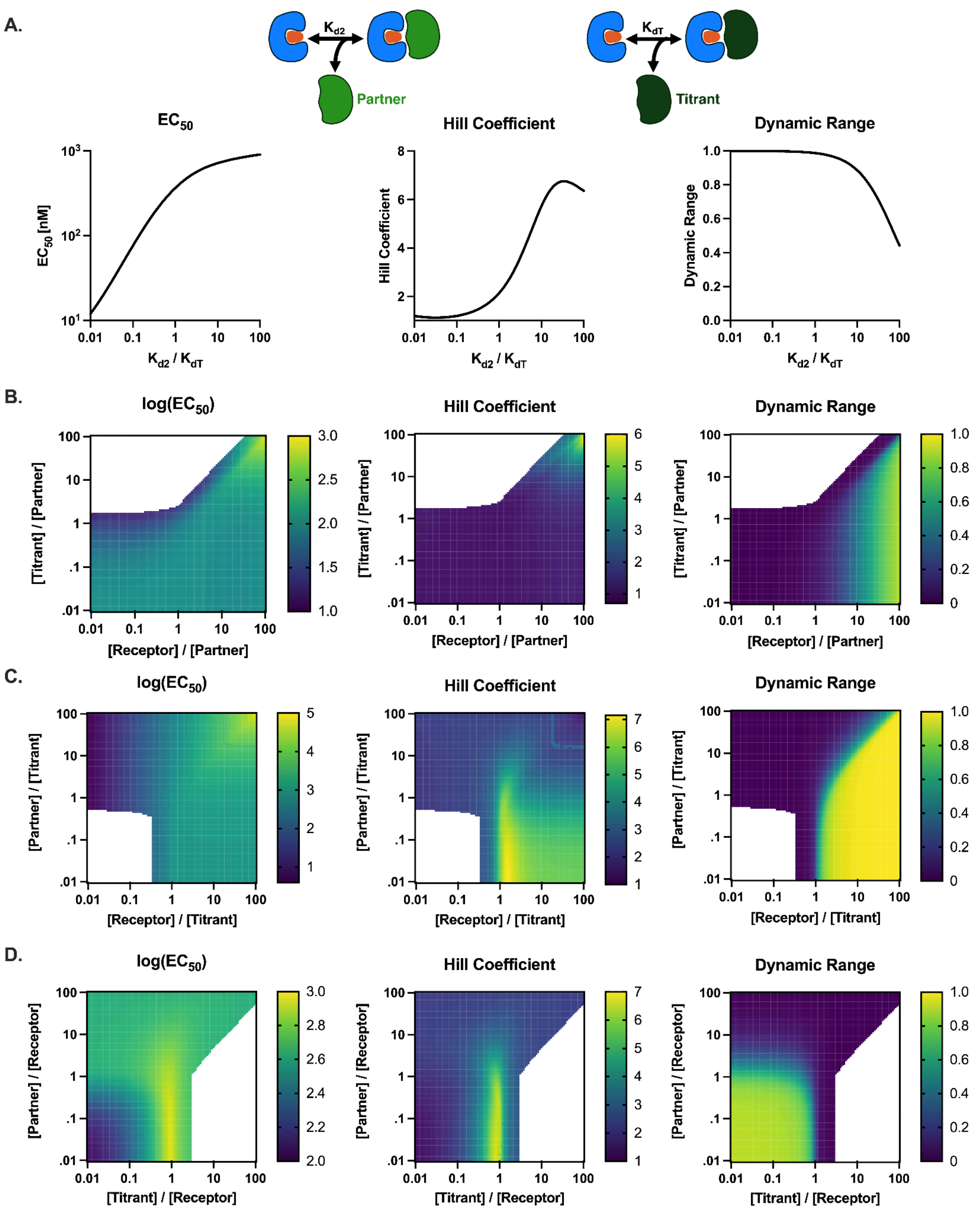


**Figure S2. Optimal phase space of the switch A.** When ligand bound receptor has higher affinity for the titrant than the partner, the $EC_{50}$ and Hill coefficient are increased however at the expense of dynamic range. Here, we fixed $K_{dT}$ at 2.1 nM and varied $K_{d2}$; 960 nM receptor, 800 nM titrant, 8 nM partner, $K_{d1}$ = 115.6 nM. **B.** When the receptor and the titrant are in great excess of the partner, the $EC_{50}$ and the Hill coefficient are maximized. Here, we fixed the concentration of partner protein at 8 nM and varied the concentrations of the receptor and the titrant. **C.** The Hill coefficient is maximized when the concentration of the receptor is in slight excess of the titrant and partner is limiting. Here, we fixed the concentration of titrant at 800 nM and varied the concentrations of the receptor and partner protein. **D.** The $EC_{50}$ and Hill coefficient are maximized when the receptor is in slight excess of the titrant and the partner is limiting. Here, we fixed the concentration of the receptor at 960 nM and varied the concentrations of the partner *and the titrant. (B-D: $K_{d1}$ = 115.6 , $K_{dT}$ = 2.1 nM, $K_{d2}$ = 105 nM).*

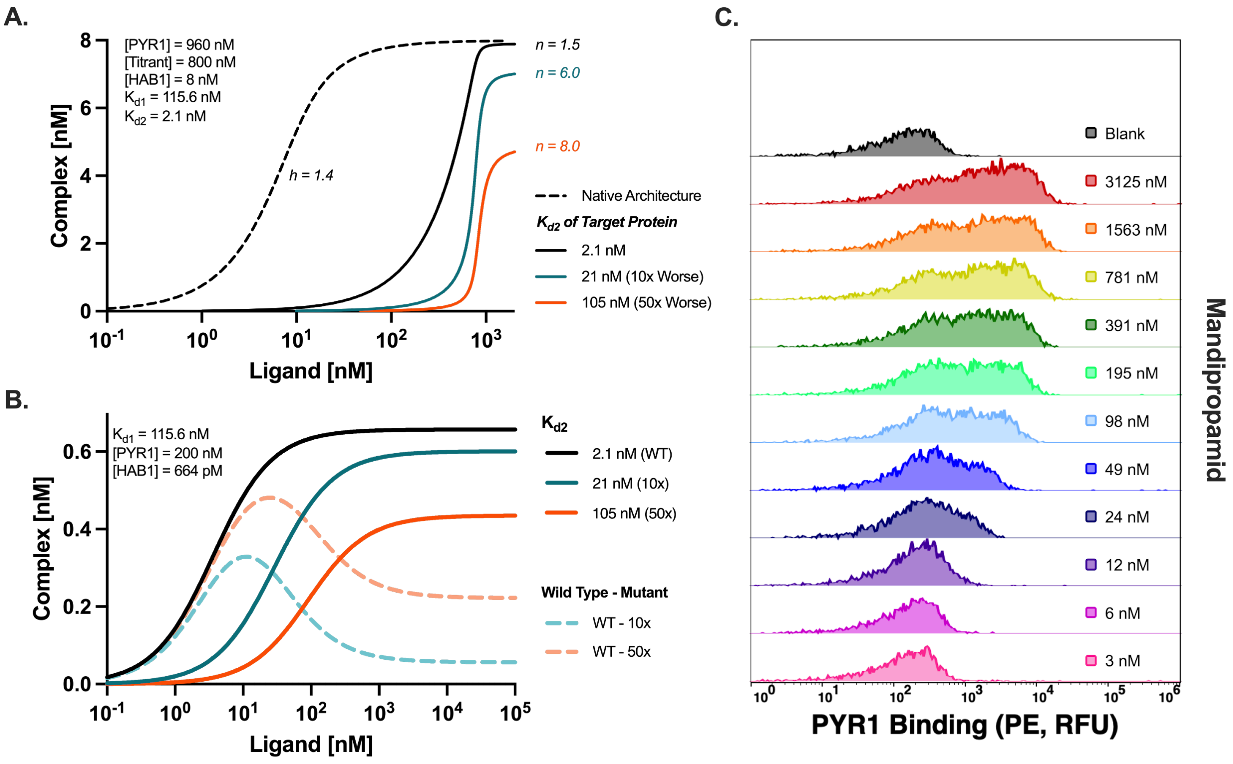


**Figure S3. Modeling informs yeast display screening conditions for the low affinity HAB1. A.** The sequestration efficacy of a titrant with $K_{dT}$ = 2.1 nM against partner HAB1s with a range of affinities. Utilizing ΔN-HAB1$^{T+}$ as the partner protein allows buffering of the system but only a minimal increase in sensitivity (n = 1.5). Partner proteins with reduced affinity can achieve much higher hill coefficients although this increase in sensitivity comes at the expense of dynamic range. A range of 10x to 50x reduced affinity was targeted in the FACS selection process to elicit a sensor enabling ultrasensitivity while maintaining dynamic range. **B.** Equilibrium binding is modeled for ΔN-HAB1$^{T+}$ and variants with 10x to 50x higher $K_{d2}$ at concentrations approximating YSD experiments. Discrimination between desired mutants and the parental sensor are shown (dotted). These results were used to inform the library YSD titration as potential concentrations to sort on. **C.** A titration was run with the HAB1 YSD library at various concentrations of mandipropamid. Histograms demonstrate the clustering and relative saturation of the library at different mandipropamid concentrations. The 391 nM and 195 nM concentrations yielded maximum dispersion between the high-affinity and low-affinity populations evidenced by the bimodal distribution. A midpoint of 290 nM was chosen to facilitate optimal discrepancy between variants during FACS selection.

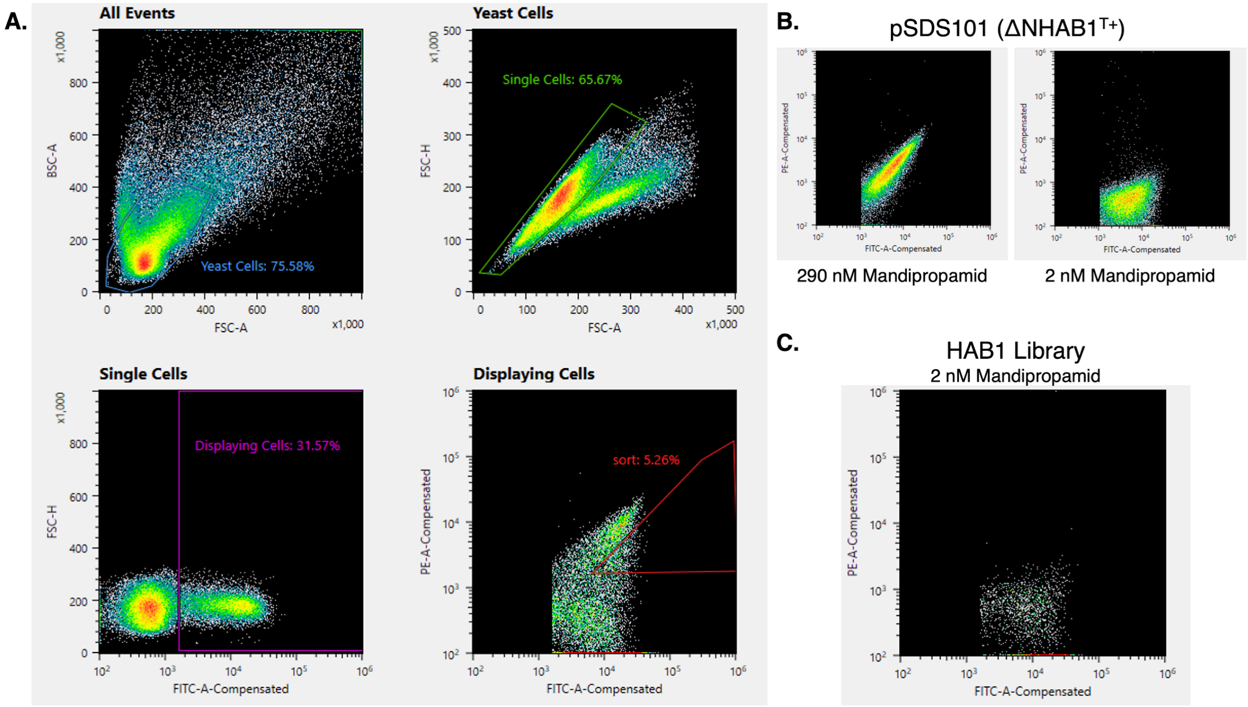


**Figure S4. Gating techniques for HAB1 library sort. A.** Forward scattering area (FSC-A) is plotted versus back scattering area (BSC-A) to gate out yeast cells. FSC-A is plotted versus forward scattering height (FSC-H) to gate out single cells. FITC is plotted on the x-axis to gate out displaying cells. PE is plotted on the y-axis and cells exhibiting an intermediate level of PYR1 binding are sorted out and collected. The yeast, single, and displaying cells gates are representative of all gating used in this study. **B.** Gated cytograms of the parental construct, pSDS101, shown as a control. **C.** Gated cytogram of the HAB1 library at 2 nM mandipropamid.

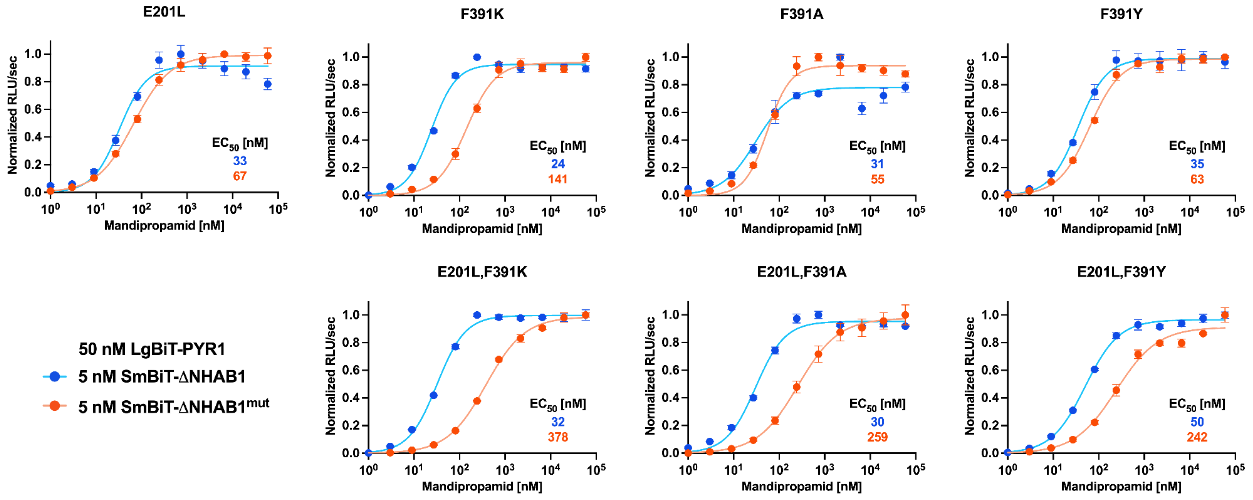


**Figure S5. *In vitro* split luciferase assay results for low-affinity HAB1 variants.** Titrations were run for each low-affinity isogenic variant (A) as well as the 3 combinatorial variants. The parental sensor was run as a control in the same plate for all setups. Data represent the mean (n = 4) and error bars represent the standard error of the mean, although in some cases the error bars are smaller than the symbols themselves. Data are fit with a nonlinear least-squares regression with a specific binding model incorporating a Hill slope.
